# *Helicobacter pylori* and Epstein-Barr virus coinfection stimulates the aggressiveness in gastric cancer through the regulation of gankyrin

**DOI:** 10.1101/2020.11.19.390807

**Authors:** Dharmendra Kashyap, Budhadev Baral, Nidhi Varshney, Anil Kumar Singh, Hem Chandra Jha

**Author notes:** Equal contributors (Budhadev Baral, Nidhi Varshney). Corresponding author **Author for correspondence:** Dr. Hem Chandra Jha, Infection Bio-engineering group, Discipline of Biosciences and Biomedical Engineering, Indian Institute of Technology Indore. Madhya Pradesh, PIN-453552, 0732-4306581.

## Abstract

Persistent coinfection of *Helicobacter pylori (H. pylori)* and Epstein-Barr virus (EBV) promotes aggressive gastric carcinoma. The molecular mechanisms underlying the aggressiveness in *H. pylori* and EBV coinfected gastric cancer is not well characterized. In the current study, we investigated the molecular mechanism involved in the cooperation of *H. pylori* and EBV-driven proliferation of gastric epithelial cells. Results showed that the coinfections are significantly more advantageous to the pathogens to create a microenvironment that favors the higher pathogen-associated gene expression. The EBV latent genes EBNA1 and EBNA3C are highly overexpressed in the coinfections compared to individual EBV infection at different time points (12 and 24 hrs). The *H. pylori*-associated genes 16s rRNA, CagA, and BabA has also been highly overexpressed in coinfections compared to *H. pylori* alone. Gankyrin is a small protein of 25 KDa involved in multiple biological and physiological processes. The upregulation of gankyrin modulates the various cell signaling pathways, leading to oncogenesis. The gankyrin shows a similar expression pattern as EBNA3C at both transcript and protein levels, suggesting a possible correlation. Further EBV and *H. pylori* create microenvironments that induce cell transformation and oncogenesis by dysregulation of the cell-cycle regulator, GC marker, cell migration, DNA response, and antiapoptotic genes in infected gastric epithelial cells by enhancing the expression of gankyrin. Our study provides new insights into the molecular mechanism where the interplay between two oncogenic agents (*H. pylori* and EBV) leads to the enhanced carcinogenic activity of gastric epithelial cells through overexpression of oncoprotein gankyrin.

**Importance:** In the present study, we have evaluated the synergistic effect of EBV and *H. pylori* infection on gastric epithelial cells in various coinfection models. These coinfection models depict the first exposures of gastric epithelial cells with EBV and then the *H. pylori.* While other coinfection models narrated the first exposures of *H. pylori* followed by the infection of EBV. This led to an enhanced oncogenic phenotype in gastric epithelial cells. We determined the coinfection of EBV and *H. pylori* enhanced the expression of oncogenic protein gankyrin. The interplay between EBV and *H. pylori* promotes the oncogenic properties of AGS cells through the newly discovered oncoprotein gankyrin. EBV and *H. pylori* mediated upregulation of gankyrin further dysregulates various cancer-associated hallmarks of genes such as cell-migratory, gastric cancer marker, tumor suppressor, DNA damage response, and proapoptotic genes.

## Introduction

Gastric cancer (GC) is the worlds’ fourth leading cause of cancer-related deaths in both males and females (1). *Helicobacter pylori (H. pylori)* and Epstein-Barr virus (EBV) are group1 carcinogens that potentially contribute to the development of GC (2). While doing so it can alter gastric physiology and immunology (3). The prevalence of *H. pylori* in about half of the global populations but it causes GC only in 3% of infected individuals (4, 5). The discrepancy in *H. pylori* infection and GC might be influenced by its strain variability such as expression of CagA, VacA, and BabA (6, 7, 8). EBV is the second most prominent cancer-associated pathogen involved in several types of lymphoid and epithelial cancers, including GC (9). Latent and lytic genes expression associated with EBV such as EBNA-1, EBNA-2, EBAN-3C, and BZLF-1 provokes the disruption of the host genes (9). The viral proteins also dysregulated the cell cycle, inflammation, angiogenesis, and tumor suppressor genes by hyper methylation activities (10, 11). Earlier studies have shown the ability of EBV for oncogenic transformation of primary gastric epithelial cells (12, 13). Thus, it is not only considered as a passive carrier but also as an active oncogenic virus contributing to early events in the development of GC (2). Lone infection of *H. pylori* or EBV is less severe in comparison to coinfection and takes more time for the development of GC (2).

Gankyrin or PSMD-10 is a recently discovered small protein of 25 KDa known to be overexpressed in various cancers (14). It is a novel oncogenic protein, involved in various cellular processes such as cell cycle progression, apoptosis, and tumorigenesis (15). Gankyrin negatively regulates the protein retinoblastoma (pRb) and tumor suppressor protein-53 (p53/TP53) to execute its oncogenic functions (16,17). Gankyrin was downregulated in GC which leads to increased cell chemosensitivity to 5-fluorouracil and cisplatin was attained by regulating cell cycle-related proteins, such as cyclin D1, cyclin E, and by activating the PI3K/AKT pathway (18).

The mutualistic relationship which endures within the microbial niche could modulate host gene expression, thus impact pathophysiology. We determined the molecular mechanism to elucidate the unexplored strategy, which involves the participation of *H. pylori* in EBV-driven proliferation of gastric epithelial cells. We have developed five different infection model including one uninfected control, using AGS human gastric epithelial cells, which is an excellent system mimicking the human gastric epithelium. We elucidated that EBV and *H. pylori* increased the gankyrin expression which led to the proliferation of infected cells in our studied system. Furthermore, we determined the status of cell migratory, cell-cycle regulatory, DNA repair, apoptosis, and tumor suppressor genes in coinfection models which can be directly linked with aggressive carcinogenesis.

## Results

### Coinfection of *H. pylori* and EBV synergistically regulates their genes expression in the gastric epithelial cell after infection establishment

In this study, we evaluated the synergistic effect of *H. pylori* and EBV on gastric epithelial cells. To understand the synergistic effect of these two pathogens we have developed the *in-vitro* coinfection models by using adenogastric carcinoma cell line (AGS). This cell line mimics the gastric epithelial cells and provides the same microenvironment for the growth of these two pathogens as the *in-vivo* conditions except pH (1.5-3.5). Before incubation of *H. pylori* with AGS cells we have checked its morphology through the Gram staining, as predicted we have observed the regular spiral shape of this bacterium **(supplementary fig.1)**. We have also observed the interaction between the *H. pylori* and EBV and how their synergistic interplay affects the pathogenic gene expression of these pathogens and downstream host cellular physiology. Our results showed that the infection-III and Infection-IV are significantly more advantageous to the pathogens to create a microenvironment that favors the higher pathogens associated gene expression as compared to individual infection at a time **(Fig2)**. The expression of pathogens associated with carcinogenic genes is multiple folds higher in coinfection models in comparison to individual infection **(Fig2 A&B)**. Both *H. pylori* and EBV synergistically provided a suitable milieu for the growth of each other. Subsequently, we have followed all the above models to understand the pathophysiology of these two pathogens in GC progression and severity and how these two pathogens mutually favor the infectivity of each other. Thus, after coinfection, we have collected the cells at various time points (12, 24, and 48 hrs) and checked the profiles of the pathogenic and host gene expression.

**Fig.1.**
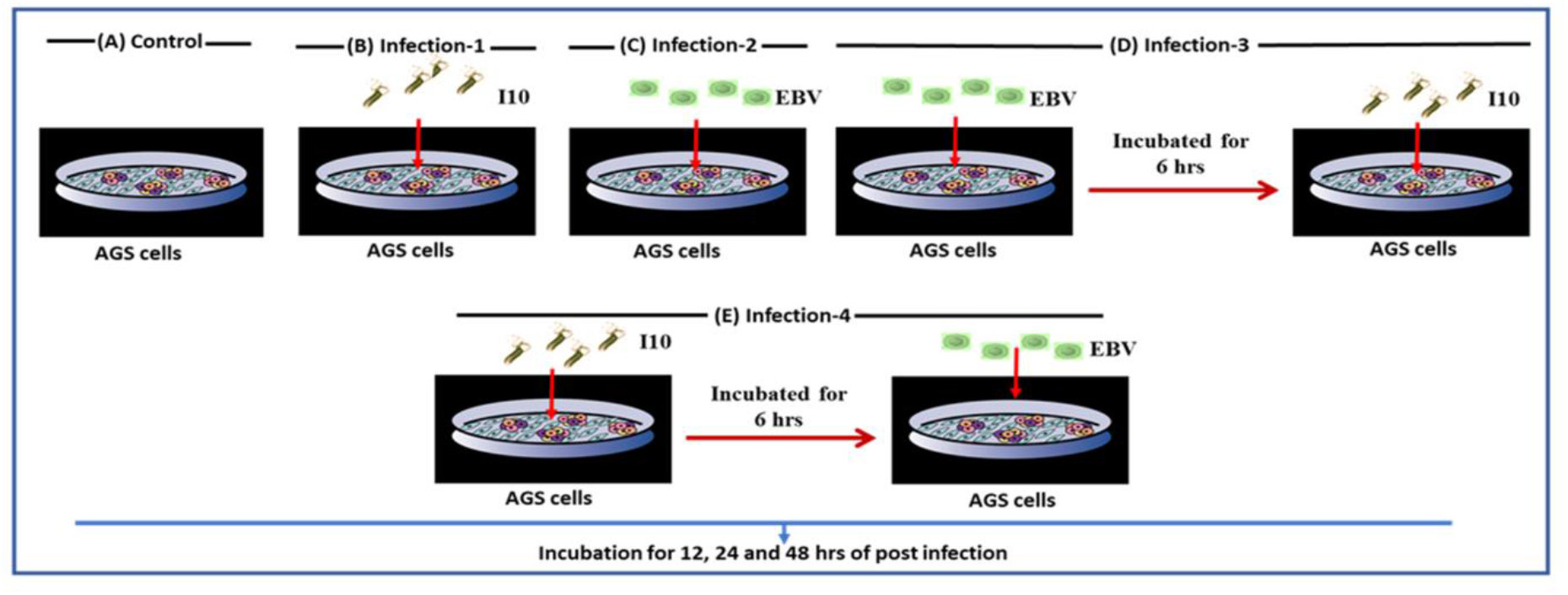
Coinfection models of *H. pylori* and EBV depicts the deep into the pathophysiology of gastric cancer. Illustration of all the studied infectious models. AGS cells were cultured in 6-well plates, and directly incubated the *H. pylori* and EBV with these cells **(A)** Uninfected AGS control cells **(B) Infection-I** portrayed the infection of *H. pylori* (I10) **(C) Infection-II** AGS were infected with EBV only. **(D) Infection-III** AGS cells were infected sequentially, first exposing the cells with EBV for 6 hrs then these exposed cells incubated with *H. pylori*. **(E) Infection-IV** AGS cells were firstly exposed with the *H. pylori* then incubated with EBV. All the infected cells were further incubated for 12, 24 and 48 hrs. These infection models were used for the whole of this study except colony formation assay.

**Fig.2.**
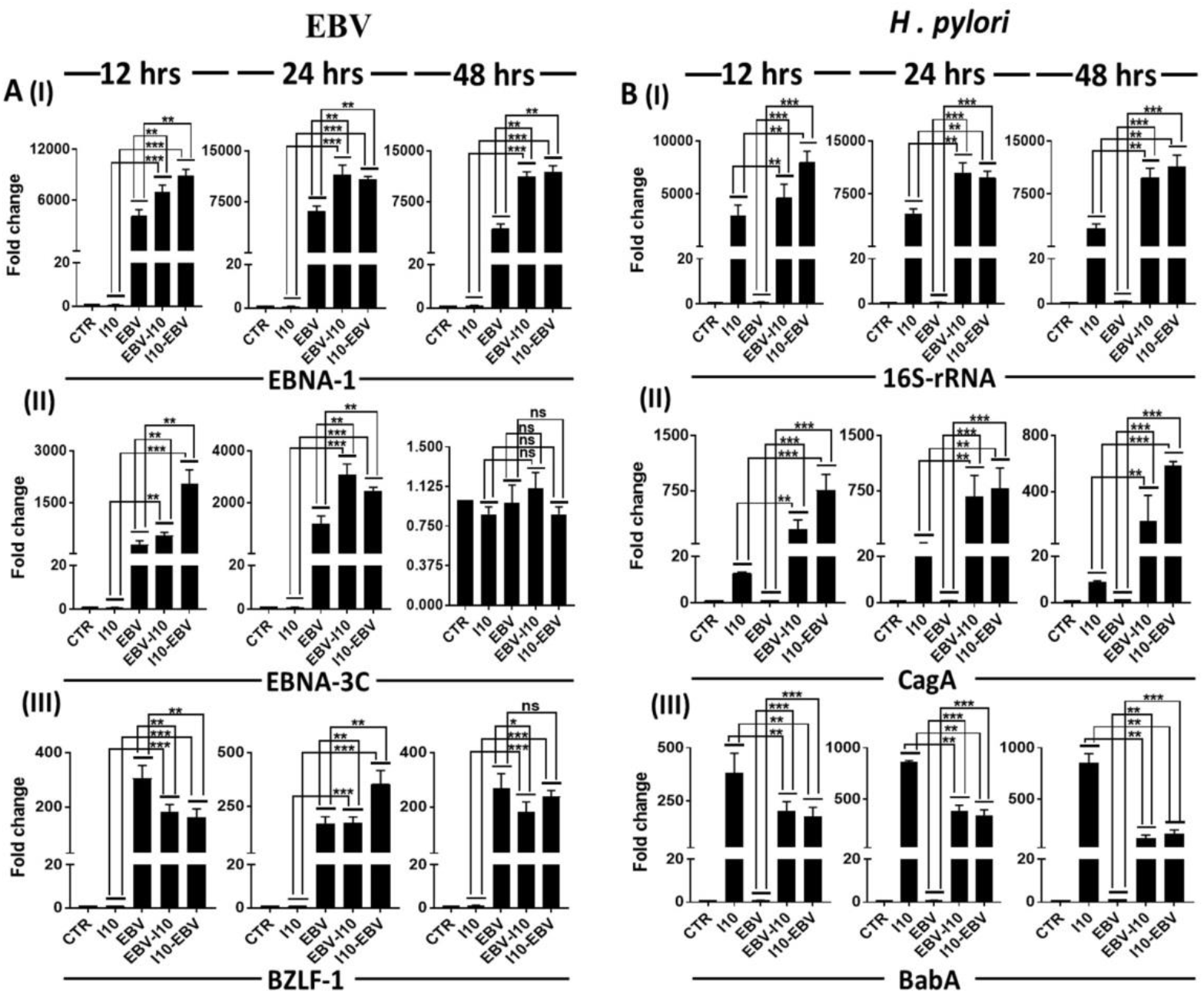
Interplay between *H. pylori* and EBV synergistically increases the transcript of its associated pathogenic genes. **(A)** Transcript expression of EBV associated latent and lytic genes. (I) & (II) Quantification of transcript of latent gene EBNA-1 & EBNA-3C respectively (III) qRT-PCR quantification of lytic gene BZLF-1 **(B)** Transcript quantification of *H. pylori* associated signature 16S-rRNA (I) and its associated pathogenic gene cytotoxin associated gene A (CagA) (II) and blood group antigen binding adhesin A (BabA) (III) in all the infection model for 12, 24 and 48 hrs post infected samples. The experiment performed for two biological and two technical replicates and the results are shown as the mean SD of two independent experiments.

After infection, we performed the qRT-PCR for the EBV-associated latent and lytic genes such as EBNA1, EBNA3C, and BZLF1 respectively. Our results showed that the expression of EBNA1 transcript is significantly higher in infection-III and IV at time point 12 (p=0.0027 and 0.054), 24 (p=0.0024 and 0.0021), and 48 (p=0.025 and 0.0021) hrs as compared to the infection-II and I **Fig2A (I)**. In this experiment, we have selected one uninfected negative control. This mock confirmed to us that there was no cross-contamination of these two pathogens and the mock was not compromised. Similarly, the nuclear antigen EBNA 3C expression was significantly higher in infection-IV at 12hrs post-infection (p=0.0450 and 0.0022) as compared to infection-II and I whereas in the case of 24hrs of coinfected samples the expression of EBNA3C was higher in infection III as compared to infection-II and I showing the interplay between the EBV and *H. pylori.* However, the expression is significantly dropped at 48hrs post-infection Fig**2A (II)**.

Furthermore, the expression of BZLF-1 transcript is higher (~100-150 folds) in infection-II, while decreasing in infection-III and IV at 12hrs post coinfection. The similar expression of BZLF-1 was determined in Infection-II and III at 24 hrs post-infection as compared to infection-IV. In 48hrs of post coinfection, there was a slightly higher expression of BZLF-1 in infection-II concerning infection-III and IV **Fig2A (III)**. Meanwhile, when comparing the infection-II, III, and IV with uninfected samples we observed that there was multiple fold higher expression of EBV associated latent and lytic genes, which confirmed the persistence of infection.

Additionally, we investigated the expression of *H. pylori*-associated genes’ transcripts such as 16s-rRNA, pathogenic gene CagA, and BabA. Our finding showed that in the case of coinfection the expression of 16s-rRNA transcripts is higher as compared to individual infection of *H. pylori* (infection-I). The expression of 16s-rRNA transcripts is higher (~2000 folds) in infection-III while it has downregulated in infection-IV at 12hrs post-infection. Moreover, the expression of this gene is about double in 24 and 48hrs as compared to 12hrs post-infection in infection-III and IV. In addition to this, the expression of 16s-rRNA transcripts increases successively from 12 (~2000 folds) to 48hrs (5000 folds) post-infection **Fig2B (I)**.

Further, we investigated the expression of well-known pathogenic genes such as CagA and BabA. Following the qRT-PCR results, we determined that the expression of CagA transcripts is significantly higher at 12 (p=0.0026 and 0.0020), 24 (p=0.0027, 0.0024), and 48 (p=0.0023 and 0.0004) hrs time point in infection-III and IV as compared to infection-I and II **Fig2B (II)**. Surprisingly, the expression of BabA is higher (~100-200 folds) at 12, 24, and 48hrs time point in infection-I while it significantly decreases in infection-II, III, and IV in all the time points such as 12, 24, and 48 hrs post-infection **Fig 2B (III)**.

### EBV and *H. pylori* regulate the expression of gankyrin in gastric epithelial cells

In addition to pathogens’ associated factors, we also determined the host factor that is regulated by the coinfection of EBV and *H. pylori.* Elevated expression of ankyrin was shown to be associated with several cancers. We investigated if this was also true for pathogen mediated gastric cancer. As expected, the infection of *H. pylori* and EBV enhanced the expression of host-associated oncogenic gene gankyrin at transcripts and protein levels. The expression of gankyrin is slightly higher at 12 and 24 hrs in infection-III and IV as compared to infection-I and II meanwhile, at 48 hrs post-infection there is significantly lower (p=0.0323, 0.0234, 0.0410 and 0.0415) expression of gankyrin in infection-III and IV whereas we have observed higher expression in infection-I and II **Fig3A (I).** Additionally, we have also observed a similar pattern of expression of gankyrin in western blot at protein level except for infection-III & IV where we examined the less expression of gankyrin protein as compared to infection-II after 12hrs of post-infection **Fig3A (II)** **and** **(III)**.

**Fig.3.**
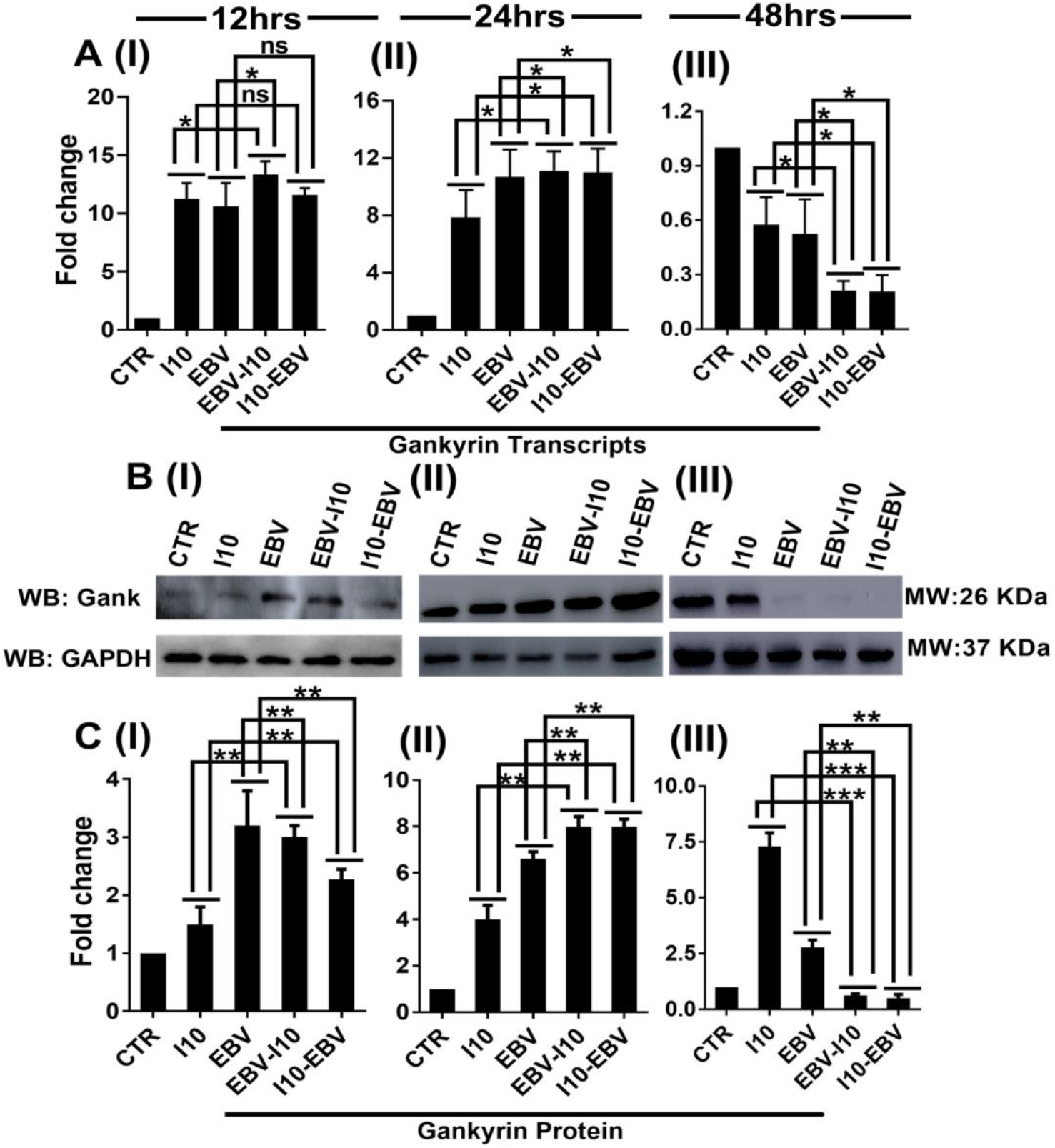
Coinfection of *H. pylori* and EBV upregulates the oncogenic protein gankyrin at both transcripts and protein levels. **(A)** Transcript expression of gankyrin in coinfection models (I), (II) & (III) depict the expression of gankyrin at 12, 24 and 48 hrs respectively. (B) (I), (II) & (III) western blot image of protein gankyrin for 12, 24 and 48 hrs post infected samples respectively. (C) (I), (II) & (III) Illustrate the quantification of blot image by Image J. software and representative graph presented in terms of fold changes for 12, 24 and 48 hrs post infected samples respectively. The significantly higher expression of gankyrin observed at 12 and 24 hrs post infected samples. While the expression of gankyrin is significantly downregulated in infection-II, III and IV in 48 hrs post infected samples. The experiment was performed for two biological and one technical replicate.

### Coinfection of EBV and *H. pylori* promotes the oncogenic properties of epithelial cells by deregulating the expression of cell-cycle regulator and the tumor suppressor genes

qRT-PCR results showed that CCND-1 gene expression is significantly higher in infection-III and IV at time point 12 (p=0.0231 and 0.0020) and 24 (p=0.0246 and 0.0349) hrs as compared to individual infected samples such as infection-I and II **Fig4A (I&II)**. Surprisingly, we witnessed the significantly lower expression of CCND-1 at time point 48hrs in the infection-II, III, and IV in comparison to infection-I (p = 0.0021) Fig4A (III). In the case of DAPK3, after 12hrs post-infection the expression of this genes’ transcripts was higher in infection-I and III as compared to infection-II and IV **Fig4B (I)** while in 24hrs post infected condition we observed the higher DAPK3 expression in infection-III and IV as compared to infection-I and II **Fig4B (II)** whereas adverse expression pattern was observed at 48hrs post-infection. Further, we have observed lower expression of DAPK3 in all infected models in contrast to uninfected control **Fig4B (III)**

**Fig.4.**
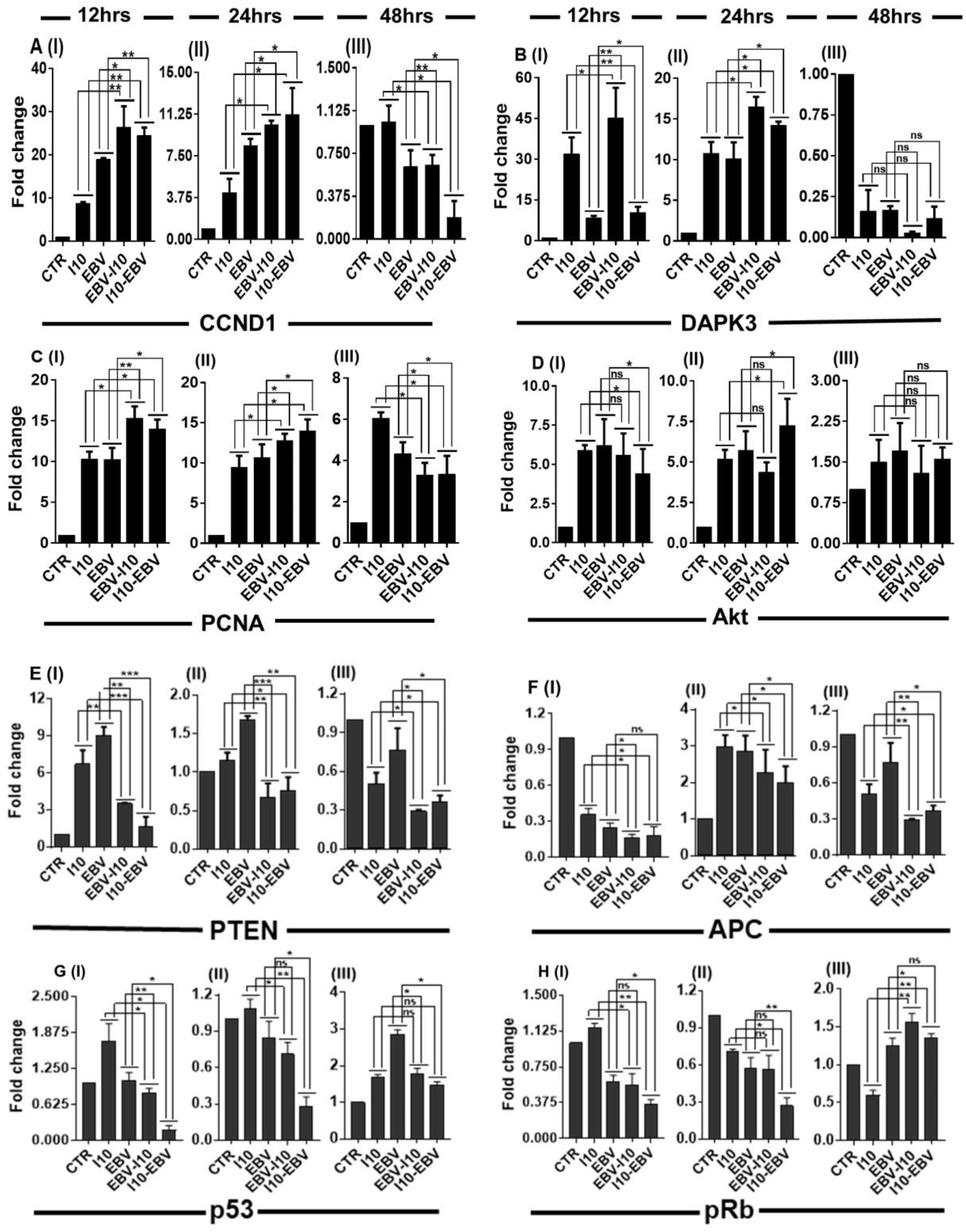
Coinfection of *H. pylori* and EBV promotes gastric cancer through the dysregulations of cell-cycle regulator and the expression of tumor suppressor gene. **(A)** I, II & III represents the cyclin D1 (CCND-1) profiles in coinfection at 12, 24 and 48 hrs respectively **(B)** I, II & III depicts the death associated protein kinase-3 (DAPK3) expression pattern at 12, 24 and 48 hrs respectively **(C)** I, II & III demonstrate the transcript expression profiles of proliferating cell nuclear antigen (PCNA) 12, 24 and 48 hrs respectively **(D)** I, II & III depicts the expression profiles of AKR thymoma (Akt) coinfected samples at 12, 24 and 48 hrs of post infection of EBV and *H. pylori*. **(E) I,** II & III illustrate the downregulation of phosphatase and tensin homolog (PTEN) in 24 and 48 hrs post infection, while it is slightly upregulated in 12 hrs post infection. **(F)** I, II & III demonstrate the expression profiles of adenomatous polyposis coli (APC) in EBV and *H. pylori* coinfected samples for 12, 24 and 48 hrs time points. **(G) & (H)** I, II & III represents the expression of p53 and pRB respectively in 12, 24 and 48 hrs post infected samples. The experiment has performed for two biological and two technical replicates and the results are shown as the mean SD of two independent experiments.

The expression of another gene PCNA was higher in 12, 24, and 48hrs post infected samples concerning uninfected control, while a lower expression of this gene was examined at 48hrs in comparison to 12 and 24hrs post infected conditions in infection-III and IV **Fig4C (I, II & III)**. The expression of another important gene Akt is comparatively higher in all infected models at 12 and 24hrs post-infection for 48hrs post infected samples **Fig4D (I, II & III)**. Further, in the case of 24hrs, we witnessed the significantly higher expression of these genes’ transcripts in infection-IV in comparison of infection-I and II (p= 0.0023, 0.0002, 0.0021 and 0.0008) **Fig4D (II)**

Tumor suppressor genes are the defense system of the body that helps to control the abnormal growth and proliferation of cells through various mechanisms such as promotes the DNA repair system, apoptosis, or modulates cell signaling. Expression of one of the important tumor suppressor gene PTEN is significantly higher in infection-I and II in comparison to infection-III and IV at 12 and 24hrs time points **Fig4E (I & II)**. The expression of this gene is downregulated in all infection models as compared to uninfected control at 48hrs of post-infection **Fig4E (III).**

Additionally, APC is another tumor suppressor gene which performs their function through the interaction of cell adhesion molecules E-cadherin and negatively regulates the expression of beta-catenin. We have evidence that the expression of APC is significantly downregulated in the case of 12hrs post-infection in all the four infection models (p= 0.0231, 0.0248, 0.0349, and 0.0423) **Fig4F (I).** Unexpectedly, the expression of APC is upregulated in 24hrs post-incubation of these two pathogens in infection-I, II, III, and IV while higher expression of this gene was observed in infection-I and II in comparison of infection-III and IV **Fig4F (II)**. Moreover, in 48hrs post infected samples, we determined the higher expression of APC in infection-II followed by infection-I and IV **Fig4F (III)**.

Furthermore, we noticed the expression of the transcripts of another important tumor suppressor gene p53 was higher in infection-I and II in comparison to infection-III and IV at 12 and 24hrs post infected conditions **Fig4G (I & II).** While in 48hrs post-infection, the expression of this gene was higher in infection-II concerning infection-I, III, and IV **Fig4G (III).**

Similarly, pRb, a tumor suppressor gene, was showing its higher expression in infection-I in contrast to infection-II, III & IV at 12 and 24 hrs post-infection **Fig4H (I & II)**. Meanwhile, the opposite expression pattern was observed in 48hrs of the infection **Fig4H (III).**

### *H. pylori* and EBV linked with the abrupt expression of gastric cancer marker and cell migratory genes by interfering with the oncogenic assessment of epithelial cells

The expression of gastric cancer marker gene Abl-1 is significantly higher in infection-III (p=0.0210) as compared to infection-I and II at 12hrs post-infection whereas we witnessed the downregulation in case of infection-IV as compared to infection-I and II **Fig5A (I).** Further, its expression is higher in all infected models in comparison to uninfected control after 24 hrs of post-co-infection **Fig5A (II)**. Surprisingly, the expression of Abl-1 is downregulated in the case of infection-II, III, and IV as compared to infection-I **Fig5A (III)**.

**Fig.5.**
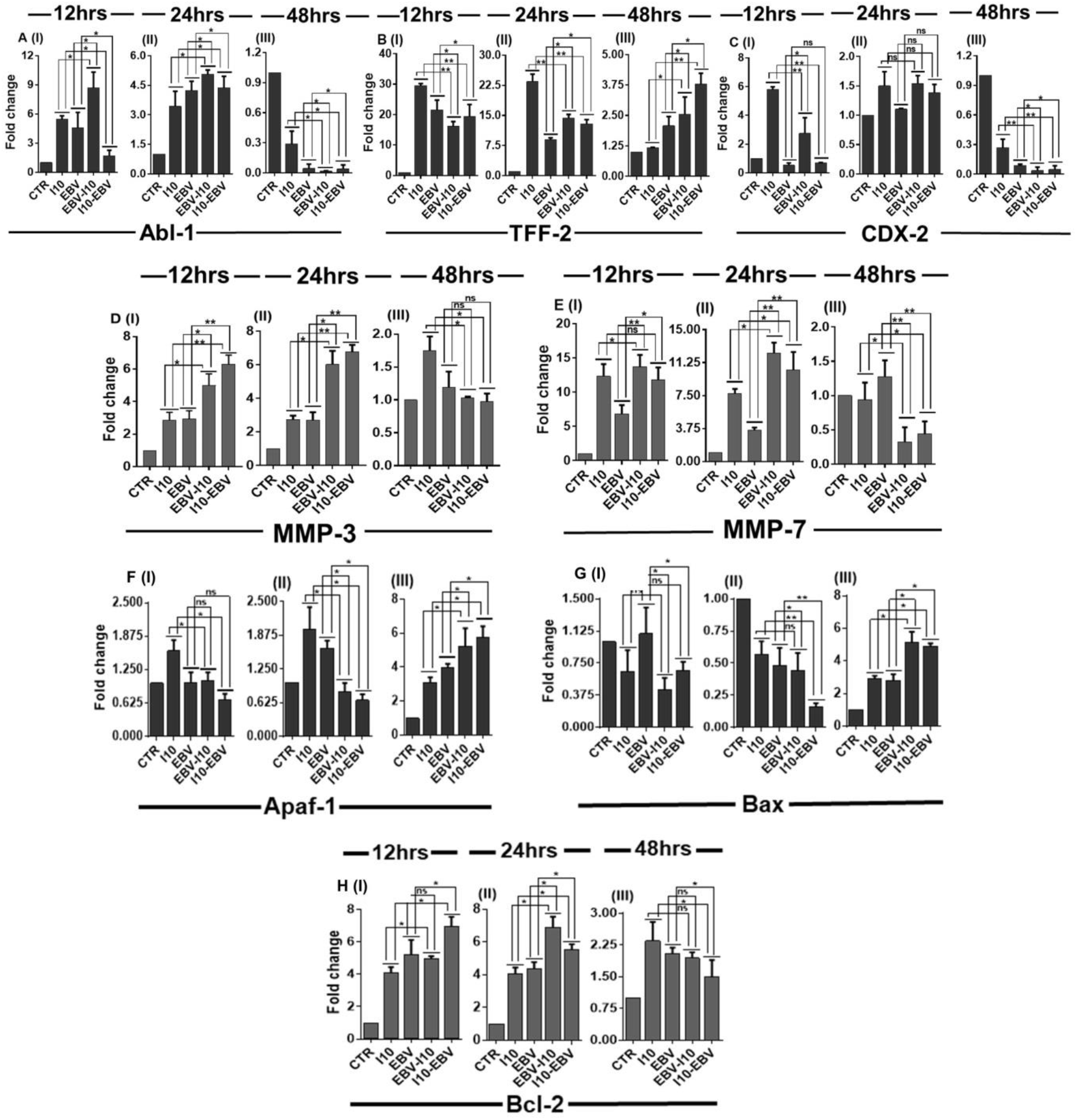
Coinfection of *H. pylori* and EBV changes oncogenic assessment of gastric epithelial cells by dysregulating gastric cancer marker, cell migratory and antiapoptotic genes. **(A)** I, II & III depicts the status of tyrosine protein kinase Abl-1 gene expression in EBV and *H. pylori* coinfection. **(B)** I, II & III demonstrate the expression profiles of trefoil factor-2 (TFF-2) at 12, 24 and hrs time points. **(C)** I, II & III illustrate the transcription factor CDX-2 in coinfected samples for 12, 24 and 48 hrs of post infection. **(D)** I, II & III depicts the expression pattern of matrix metalloprotease-3 (MMP-3) in 12, 24 and 48 hrs post infected samples. **(E)** I, II & III represents the expression status of matrix metalloprotease-7 (MMP-7) **(F)** I, II & III illustrate the downregulation of Apaf-1 **(G)** I, II & III represents the decrease in amount of Bax transcripts in coinfected samples at 12, 24 and 48 hrs of post infected samples. **(H)** I, II & III represents the expression pattern of BCL-2 in which at 12 and 24 hrs post infection the expression of antiapoptotic gene is elevated as the expression of gankyrin is also several folds upregulated in 12 and 24 hrs post infected samples in above mentioned results. While the expression of BCL-2 follows the same pattern at 48hrs post infected as the expression of gankyrin was observed. The experiment has performed for two biological and two technical replicates and the results are shown as the mean SD of two independent experiments.

Adversely, the higher expression of TFF-2 was observed in infection-I and II in comparison with uninfected control and infection-III and IV **Fig5B (I)**. In the case of 24hrs post-infection, we determined the significantly higher expression of TFF-2 in infection-I followed by the infection-III and IV **Fig 5B (II)**. Besides these, in 48hrs post, infected samples the expression of TFF-2 increased successively from infection-I to IV while the expression of this gene is comparatively alleviated in respect to 12 and 24hrs of post-infection **Fig5B (III)**. Expression of another gastric cancer marker gene CDX-2 is higher in infection-I and III in comparison to infection-II and III at 12hrs of post infected samples **Fig5C (I)**. CDX-2 expression was slightly higher in infection-I as compared to other infection models in 24hrs post infected samples **Fig5C (II)**. In addition to the above results it was significantly decreased in infection-I, II, III, and IV in 48hrs post-infected samples **Fig5C (III)**.

The upregulation of cell migratory genes is a hallmark for cancer cell metastasis. We witnessed the expression of MMP3, another cell migratory gene that helps the cells to metastasize, was also significantly upregulated in infection-III and IV in comparison of infection-I and II in 12, and 24hrs post infected samples **Fig5D (I & II)**. Additionally, after 48hrs post-infection of EBV and *H. pylori* the expression of MMP3 is moderately higher in infection-I while in infection-II, III, and IV the expression of these gene transcripts relatively similar to uninfected control **Fig5D (III).** Although the results revealed that expression of MMP7 is upregulated in infection-III and IV in contrast to infection-I and II at 12 and 24hrs of post-infection **Fig5E (I, II),** while we observed the adverse expression in 48hrs of post-infection **Fig5E (III)**.

### EBV and *H. pylori*-associated pathogenic gene expression could cause GC by regulating DNA damage response genes and anti-apoptotic gene Bcl-2

The effective function of DNA damage response genes in humans is to provide stability and maintain its genome integrity. Moreover, cancer-causing infectious agents such as EBV and *H. pylori* dysregulated the status of DNA damage response transcripts by changing the milieu of the surrounding gastric epithelial cells. Interestingly, we determined the expression of DNA damage response gene Apaf-1 expression is slightly higher in infection-I in 12hrs post infected samples. A quite similar expression was observed in infection-II, III, and IV in comparison to uninfected control **Fig5F (I)**. Furthermore, the expression of this gene at 24hrs is similar to the 12 hrs post infected samples in all the studied models except infection II where we observed higher expression of Apaf-1 **Fig5F (II).** Meanwhile, the expression of Apaf-1 in 48hrs post infected samples successively increases from the infection-I to IV **Fig5F (III).**

In addition to Apaf-1, the expression of Bax is significantly downregulated in 12hrs post infected samples in all the conditions except in infection-II **Fig5G (I).** While the expression of Bax is successively downregulated in all the four infections in 24 hrs post infected samples **Fig5G (II)**. Furthermore, this gene showed higher expression in infection-I, II, III, and IV in comparison of uninfected control in 48hrs time point, whereas, in infection-III and IV the expression of Apaf-1 was about 1.5-fold higher than the infection-I, II **Fig5G (III)**.

DNA damage response genes perform their function through its anti-apoptotic properties hence, the overexpression of anti-apoptotic genes such as Bcl-2 helps in cell survival and proliferation. Pathogens such as EBV and *H. pylori* modulate the expression of this type of host protein and promote the severity of gastric cancer. Coinfection of EBV and *H. pylori* upregulates the expression of anti-apoptotic gene Bcl-2 in 12hrs post-infection in all the four infection models compared to uninfected control whereas, we determined significantly higher expression of this gene in infection-IV **Fig5H (I)**. In addition to 12hrs, the similar expression of Bcl-2 is observed in 24hrs post infected samples except for the infection-III in which we have detected the higher expression of Bcl-2 **Fig5H (II)**. While in 48hrs post infected samples the expression of Bcl-2 is still higher compared to uninfected control. Moreover, the expression of this gene successively decreases from the infection-I to IV **Fig 5H (III)**.

### Coinfection of EBV and *H. pylori* promotes GC through the upregulation of oncogenic protein gankyrin in a time-dependent manner

Coinfection of EBV and *H. pylori* enhances the expression of oncogenic protein gankyrin in a time-dependent manner. As in the above-mentioned results, we revealed that the EBV and *H. pylori* could cause gastric cancer through the upregulation of gankyrin transcripts. To further support our qRT-PCR results we performed the immunostaining of gankyrin after coinfection of EBV and *H. pylori* by following the above-mentioned infection models (infection-I, II, III, and IV). Surprisingly, we obtained the similar expression pattern of gankyrin in immunostaining as observed at the transcript level. While, the higher expression of protein gankyrin is observed in all the four infectious models moreover, in infection-III and IV we evident that there is a significantly higher expression of gankyrin in 12hrs post-infected samples **Fig6A (I&II)**. Interestingly, in 24hrs post infected samples the expression of gankyrin remains higher in infection-III and IV in comparison of infection-I and II **Fig6B (I&II).** Furthermore, we determined the significantly downregulated expression of gankyrin at 48hrs post infected samples in all four studied models in comparison to 12 and 24hrs post infected samples **Fig6C (I&II)**.

**Fig.6.**
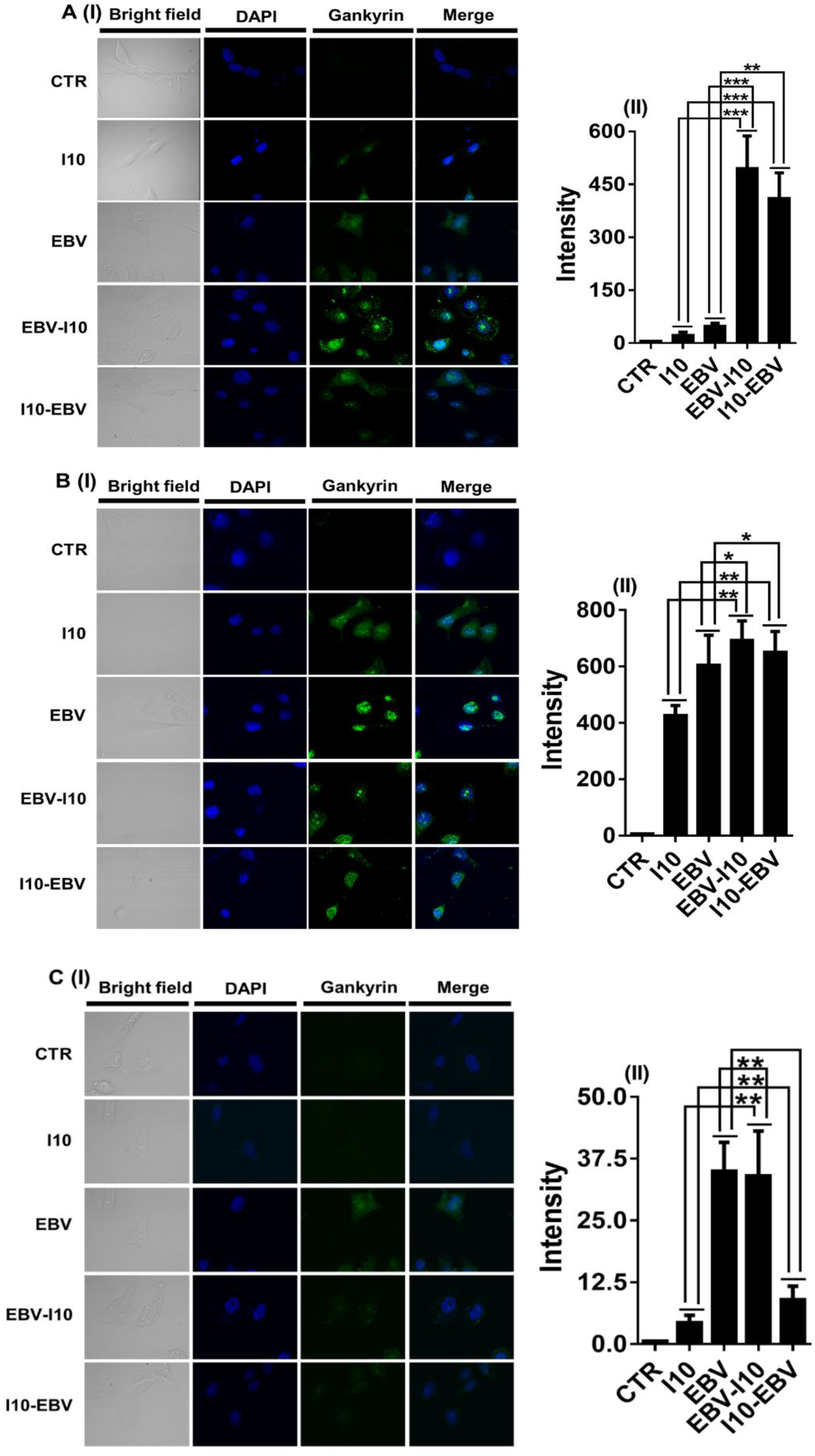
Coinfection of EBV and *H. pylori* causes gastric cancer through the upregulation of oncogenic protein gankyrin. Immunofluorescence results of oncogenic gene gankyrin validate our previous transcripts and western blot results. Graphical representation of quantified immunoblotting image through the Image J. software. After coinfection of EBV and *H. pylori* we observed elevated protein expression of gankyrin in all the time points for all studied infection except 48 hrs post infection in which observed downregulation of gankyrin in infection-II, III and IV. **(A) I, (B) I & (C) I** Representative immunoblotting image of gankyrin represents the protein expression of gankyrin after 12, 24 and 48 hrs of post infection respectively. Graphical representation of quantified immunoblotting image through the Image J. software. First panel showed an uninfected mock. Second and third panel illustrate the gankyrin protein after I10 and EBV infection. While the fourth and fifth panel shows the intensity of gankyrin expression after EBV-I10 and I10-EBV post infection. **(A) II** Approximately 600-700-fold increased expression of gankyrin observed in EBV-I10 and I10-EBV coinfected samples as compared to individual infection of I10 and EBV after 12 hrs post infection. **B (II)** The expression pattern of gankyrin in 24 hrs post infected samples is about 100 to 200-fold higher in EBV-I10 and I10-EBV coinfection. **C (III)** The expression of oncogenic protein gankyrin after 48 hrs of post infection is about 3-32 higher in infection III and IV in comparison of infection-I. The expression of gankyrin in infection-II is about equal to infection III while the 25-30-fold lower expression of gankyrin was observed in infection IV. The experiment has performed for two biological and three technical replicates and the results are shown as the mean SD of two independent experiments.

### Endogenous and exogenous expression of gankyrin promotes the cell proliferation by dysregulating the cell migration, cell-cycle regulator, protooncogene, gastric cancer marker, and tumor suppressor gene in AGS cells

Protein blot image showed the exogenous gradient expression of gankyrin in AGS cells after 48hrs of post-transfection of the plasmid containing gankyrin gene **Fig7 (A)** **and** **B**. The result revealed a concentration-dependent increase in gankyrin expression. Moreover, to support our hypothesis that EBV and *H. pylori* cause gastric cancer via the upregulation of gankyrin we exogenously overexpressed gankyrin in AGS cells and performed cell proliferation assay. Surprisingly, results showed that the exogenous expression of gankyrin promotes cell proliferation **Fig7 (C).** Furthermore, to validate our previous results the coinfection of EBV and *H. pylori* upregulated the expression of oncogenic protein gankyrin which in turn changes the cell microenvironment and promotes cell proliferation we performed the cell proliferation assay by following all the four studied infection models. Interestingly, we observed that the rate of cell proliferation is comparatively significantly higher than the uninfected mock. Besides this in 12, 24, 36, and 48hrs post infected samples the number of viable cells is higher in infection-II, III, and IV in comparison to infection-I and mock **Fig7 (D)**

**Fig.7.**
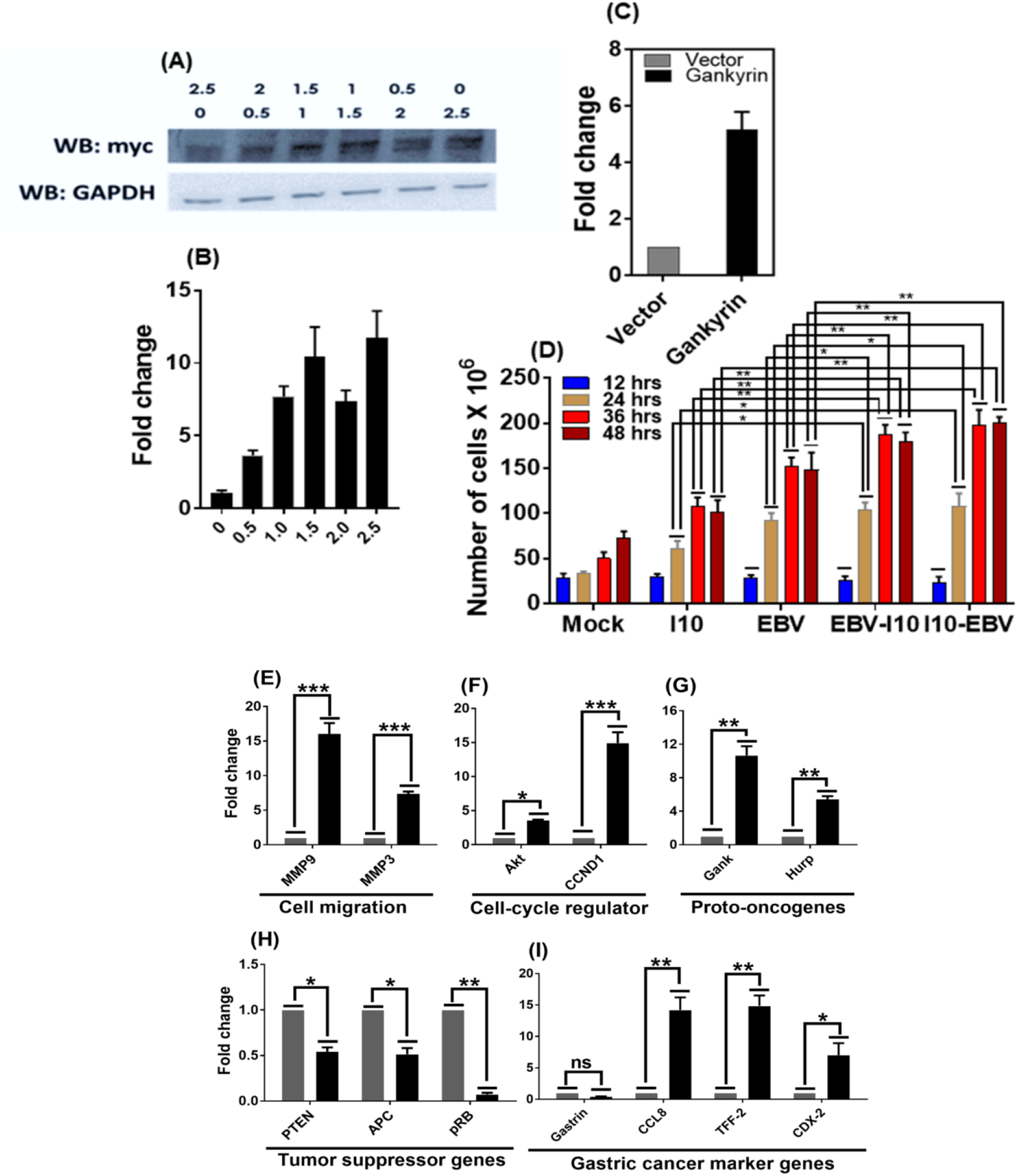
Coinfection mediated upregulation of gankyrin and ectopic expression of gankyrin promotes the rate of cell proliferation by interfering with the expression of various cellular genes. **(A)** Western blot image represents the ectopic expression of oncogenic protein gankyrin. **(B)** Graphical representation of increased concentration gradient of gankyrin quantified by the image J software of western blot image. **(C)** Shows the ectopic expression of gankyrin enhances the cell proliferation in trypan blue cell exclusion method of cell counting **(D)** Moreover, coinfection of EBV and *H. Pylori* also enhances the rate of cell proliferation in coinfected samples and may be linked with the increased expression of gankyrin. **(E)** Represents the ectopic expression of gankyrin significantly enhanced the expression of cell migratory gene MMP-7 and MMP-3. **(F) & (G)** Representative graph shows the elevated expression of cell-cycle regulatory gene Akt and CCND-1 and proto-oncogenes hepatoma upregulated protein (Hurp) respectively. Moreover, overexpression of gankyrin significantly alleviates the expression of tumor suppressor gene PTEN, APC and pRB shown in **(H).** While the expression of gastric cancer marker gene gastrin expression is slightly lower followed by the upregulation of C-C motif chemokine ligand-8 (CCL-8), TFF-2 and CDX-2 **(I).** The experiment has performed for two biological and two technical replicates and the results are shown as the mean SD of two independent experiments.

Interestingly, exogenous overexpression of gankyrin results showed that the oncogene gankyrin play their role via the interaction of various host cellular physiological pathway as it acts differentially on a various cellular mechanism such as cell migration, cell-cycle regulator, proto-oncogenes, gastric cancer marker, and tumor suppressor genes. Exogenous overexpression of gankyrin significantly upregulated the cell migratory gene MMP7 and MMP3 **Fig7 (E)** and the cell cycle regulator transcripts of Akt and CCND-1 **Fig7 (F)**. **Fig7 (G)** Showed the ectopic expression of gankyrin which enhances the expression of Hurp. Interestingly, the expression of tumor suppressor genes PTEN, APC, and pRB significantly downregulated in exogenously overexpressed samples **Fig7 (H)**. Furthermore, the ectopic overexpression of gankyrin increases the expression of gastric cancer marker gene CCL8, TFF-2, and CDX-2 while it downregulated the expression gastrin **Fig7 (I)**.

### Coinfection of EBV and *H. pylori* promotes the cell migration linked with the gankyrin

We further investigated whether the cellular physiological changes elicited by EBV and *H. pylori* could contribute to cell migration using a wound-healing assay. This assay demonstrated that EBV and *H. pylori* stimulated motility in the AGS cells. As expected, we observed the higher cell migration in infection-III and IV (p = 0.0025, 0.0023, 0.0022, and 0.0027) **Fig8A (I)** in all the selected time points. Furthermore, we counted the migratory cell and represented in graph **Fig8A (II)**

**Fig.8.**
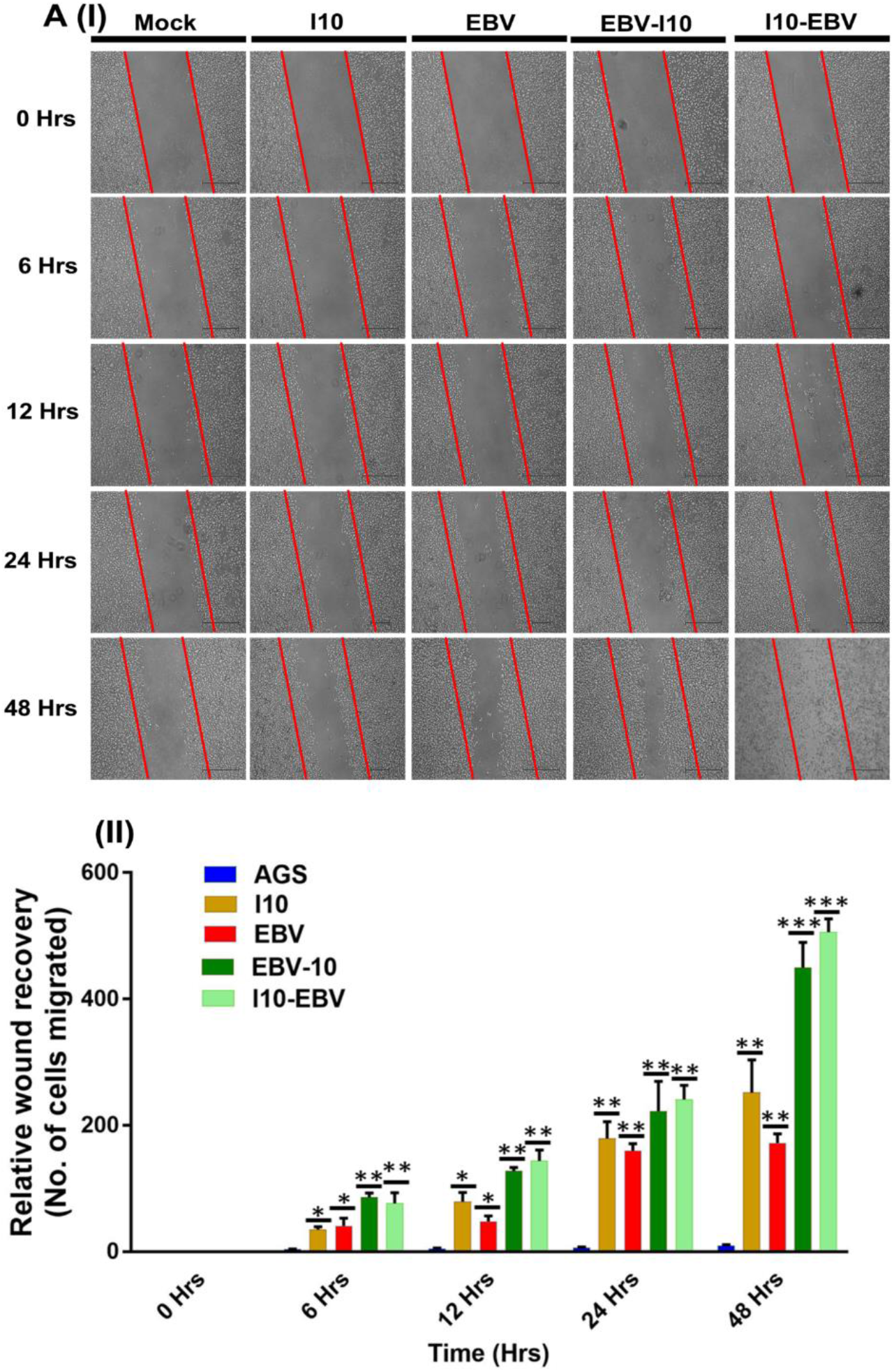
Coinfection of EBV and *H. pylori* enhanced the efficiency of cell migration of AGS cells. **(A) I** Representative image of scratch wound healing assay. First panel shows the uninfected AGS cells. Second, third, fourth and fifth panel shows the infection-I, II, III and IV respectively for the 0, 6, 12, 24 and 48 hrs of post infection. **(A) II** Graphical representation of the number of cells migrated in each infection and uninfected mock.

### Coinfection and ectopic overexpression of gankyrin promotes the oncogenic properties of AGS cells

We performed the clonogenic assay to select the EBV positive cells for the understanding of oncogenic transformation efficiency of coinfected samples in comparison to individual infection. The pore size of the transwell is smaller than the *H. pylori* only allowed the *H. pylori* secretory molecules to reach the AGS cells, not the *H. pylori*. The experimental models are depicted in **Fig9 (A)**. The lower number of colonies was observed in uninfected and *H. pylori*-infected AGS cells. Besides these the density of colonies is more in EBV infected AGS cells followed by the significantly higher number of colonies were determined in coinfected samples **(B).** Further, we quantified these colony numbers into the density and observed that the density of colonies was higher in EBV and *H. pylori* coinfected samples **(C)**. As we speculated, we observed that the secretory molecules which are secreted by *H. pylori* may be linked with the increased EBV virion in AGS cells. To further validate our finding the coinfection of EBV and *H. pylori* promote the oncogenic properties of AGS cells through the upregulation of gankyrin. We exogenously overexpressed the gankyrin in AGS cells and determined that the ectopic expression of gankyrin enhanced the clonogenic properties of AGS cells **(D).** We quantified the densities of colonies and represented them in graph **(E)**.

**Fig.9.**
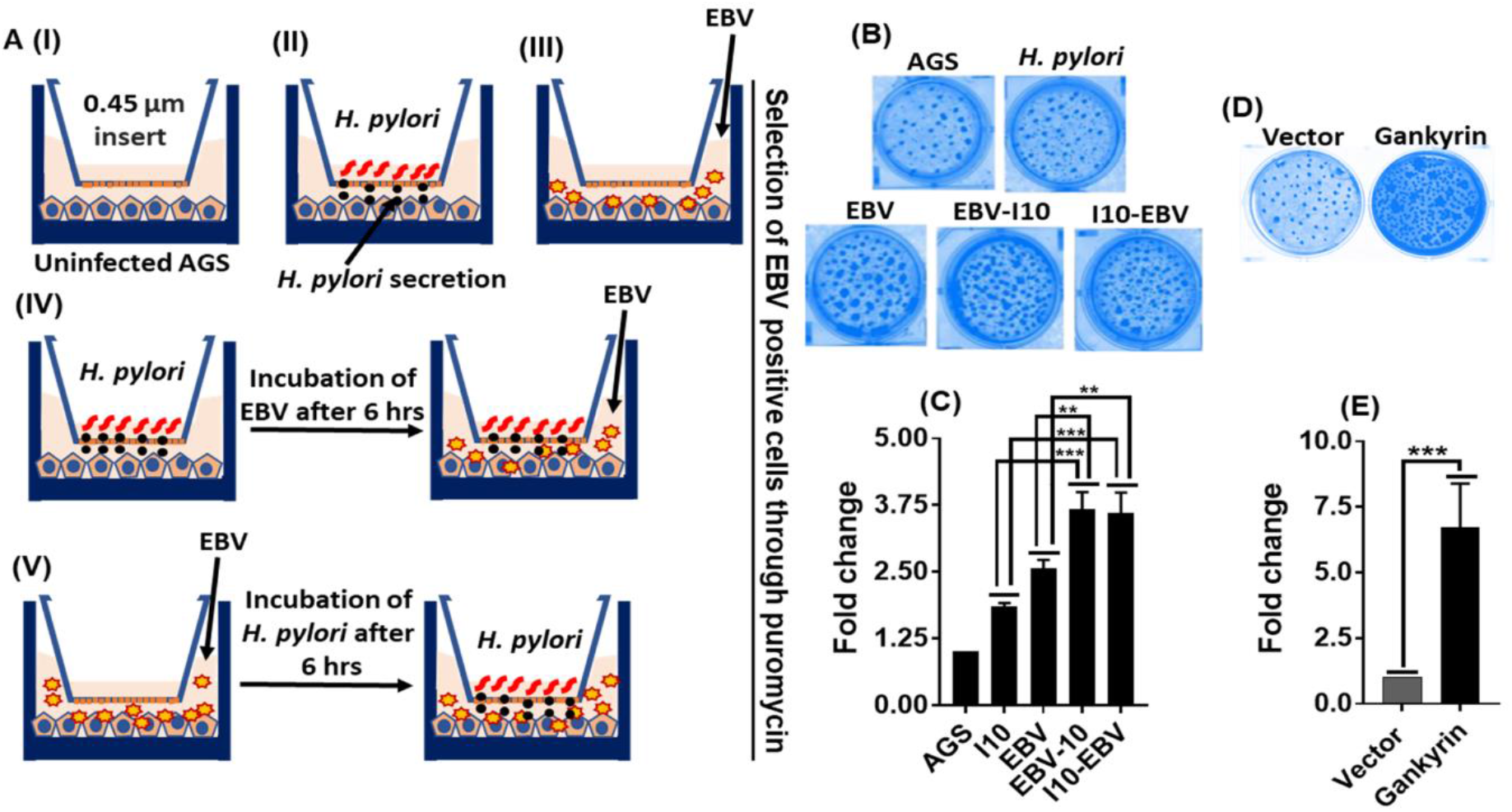
Coinfection and ectopic overexpression of gankyrin can be linked with the elevated oncogenic properties of AGS cells. **A (I)** Uninfected AGS cells. **(II)** AGS cells with transwell and *H. pylori* infection. **(III)** AGS cell only with EBV infection. **(IV)** First infection of *H. pylori* for 6 hrs followed by second infection of EBV. **(V)** First infection of EBV for 6 hrs followed by second infection of *H. pylori*. Selection of EBV positive colonies for 14 days. **(B)** Representative image of foci formation after coinfection. **(C)** Quantification and graphical representation of density of foci through the Image J. software. **(D)** Further we have validated foci formation results through the ectopic expression of gankyrin in AGS cells and selected the gankyrin positive cells through the geneticin (G-418) for 14 days. Representative image of foci in exogenously overexpressed AGS cells. **(E)** Quantification and graphical representation of density of foci in exogenously overexpressed AGS cells through the Image J. software. The experiment has been performed three times and the results are shown as the mean SD of two independent experiments.

## Discussion

Chronic or acute infection of pathogens that destroy the immune system is a severe threat to human health and can cause diseases like Alzheimer’s, meningitis, cancer, etc (19–21). Infectious agents not only induced the carcinogenesis but it is also involved in the promotion of its aggressiveness (22, 23). About one-fifth of total human oncogenesis is associated with infectious agents. Pathogens modulate the cellular milieu by dysregulation the gene expression, fit for the potential carcinogenesis (24, 25). Studies suggested that *H. pylori* and EBV promote oncogenesis by reframing the expression of cellular genes at the level of transcription and translation in infected cells (2). Reprogramming of expression of transcript and protein could promote the intrusiveness of these lesions. In this study, AGS cell lines were used which mimic the primary gastric epithelial cells except for adherent’s junction. It is well established that *H. pylori* and EBV infect the AGS cells and NCI-N87 in equal efficiency (2, 25). Thus, all these above reasons elicited us to investigate the synergistic effect of *H. pylori* exposed AGS cells for the successful infection of EBV and further how the microenvironment of EBV infected AGS cells favour the fortunate infection of *H. pylori*.

Various studies reported that *H. pylori* exposed cellular environmental conditions are suitable for the enhanced EBV infection and vice-versa (2, 25). Moreover, we also observed the enhanced EBV-GFP fluorescence in fluorolog analysis at various time intervals in all the coinfection models except in infection-I (data not shown). In a study by Pandey et. al. and Sonkar et. al, reported that these pathogens dysregulated the cellular genes’ transcript expression which led to the aggressive GC (2). Moreover, till yet it is an enigma whether the prior exposures of *H. pylori* create a suitable niche for the EBV infection or anterior windage of EBV creates a niche that favors the aggressive growth of *H. pylori* and further gastric carcinoma. Interestingly, we got similar results as being speculated, coinfection of EBV and *H. pylori* enhanced the pathogenic gene expression such as EBNA-1, EBNA3C, and BZLF-1 in various time intervals. Various studies reported that EBNA-1 and EBNA-3C enhanced the G1/S phase transition, cell invasiveness, and metastasis through the upregulation of cell signaling genes such as slug, snail, vimentin, and E-cadherin (23, 26–29). In addition to these functions, it is also known to regulate tumor suppressor genes like p53, pRb which play an important role in oncogenesis and cancer progression.

In this study, we determined the expression of EBV associated latent genes (EBNA-1 and EBNA3C) can significantly promote the replication and spread of the virus in coinfected conditions up to 48hrs then the individual infection. The gene expression pattern in *H. pylori* and EBV coinfections revealed that *H. pylori* contributed to the increased replication and transcription of EBV pathogenic genes and virion production. Surprisingly, we observed that another latent gene EBNA3C expression is significantly correlated with the expression of oncogenic protein gankyrin at 12, 24, and 48hrs post infected samples. To support this finding, we performed the western blot analysis in which we observed the same expression pattern of gankyrin as we observed in qRT-PCR. These results revealed that the EBV-associated EBNA3C may be directly linked with the expression of host oncogenic protein gankyrin as the expression of EBNA3C and gankyrin was alleviated in the only presence of EBV. The current study suggested that EBNA3C could directly regulate the host oncogenic protein gankyrin expression. These results supported that EBNA3C may cause gastric cancer through the regulation of host oncogenic protein gankyrin. It is an established oncoprotein which is found to be overexpressed in various cancers and involved in various biological processes, encompassing transformations of normal cells into cancerous one (15). Abnormal expression of gankyrin observed in human liver cancer, pancreatic cancer, oesophageal cancer, cervical, lung cancer, breast cancer, and glioma (16). Although there is no report available till which suggests that infectious agents caused the GC through the upregulation of oncogene gankyrin?

This is the first study demonstrating that the exposure of AGS cells with EBV before *H. pylori* infection and vice-versa increased the expression of gankyrin which led to aggressive gastric carcinogenesis. The expression pattern of bacterial signature gene 16s-rRNA indicated the establishment of infection in AGS cells. *H. pylori*-associated secretory antigen CagA encoded by the CagA-pathogenicity island (CagA-PAI) present in about 89% of *H. pylori* strains caused morphological dysplastic alteration in epithelial cells (30), (31). Dysplastic changes are important and contribute to the initiation of carcinogenesis. Furthermore, *H. pylori* encoding CagA is associated with hustle inflammation, peptic ulcers, gastritis, and GC (24).

In the present study, we determined that the human gastric epithelial cells have enhanced susceptibility to EBV infection in the presence of CagA positive *H. pylori* strain (I10). Pandey *et al.* showed that a CagA mutant strain of *H. pylori* has a significant reduction in viral titer production (2). This study was suggested that instead of CagA mutant, wild type *H. pylori* contributed significantly to viral replication and subsequently cancer progression and aggressiveness. *H. pylori* loaded with type-IV bacterial secretion system that helps in the injection of its associated secretory protein into the host-derived cells. Alternative mode of uptake of CagA through integrins mediated rearrangements of actin filaments and further endocytosis. Endocytosis of CagA further changed the intracellular milieu and led to the morphological alteration in the gastric epithelial cells. Here, our results supported the hypothesis that the type-IV secretory mechanism was used by this bacterium, as the direct incubation of wild type *H. pylori* showed enhanced infectivity. The expression of signature sequence 16s-rRNA, CagA were higher in coinfected samples which concluded that the virus could also help synergistically to increase the expression of *H. pylori* associated factors. Expression of 16s-rRNA inferences that enhanced replication and multiplication of *H. pylori* genome in presence of EBV. BabA transcripts are known to be highly expressed in peptic ulcer and gastric cancer rather than asymptomatic colonization. Similar observations were made with the BabA transcript in infection-I which showed that the expression of BabA is important for enhanced infectivity. Further, the higher BabA expression early point suggested the increased infectivity of *H. pylori* as it helps in the attachment of *H. pylori* with host cells.

In addition to the above results, the recruitment of gankyrin by the coinfection dysregulated the downstream genes which were known to be directly influenced by upregulation of this protein. These genes are studied under six categories such as cell cycle regulator genes, GC marker, cell migratory, tumor suppressor, DNA damage response, and antiapoptotic gene. Moreover, we investigated that EBNA-3C and gankyrin can further change the status of various cellular proteins which have played a significant role in gastric carcinogenesis. Although, experiments need to be conducted to validate the one-to-one relation of EBNA3C and gankyrin. Results of the present study suggested that the differential expression of cell cycle regulatory genes such as CCND-1, DAPK3, PCNA, and Akt in all the infectious models at various time points could be linked with the increased oncogenesis. Gankyrin upregulated the CCND-1 and PCNA in pancreatic and gastric cancer respectively (32). Besides these genes, the expression of gankyrin upregulates the Akt at transcript and protein level in hepatocellular, liposarcoma, and gastric cancer (33–35). The upregulation of Akt protein through the action of gankyrin enhanced cell proliferation, metastasis, stem cell self-renewal capacity, autophagy, and chemoresistance (18, 36). The upregulation of death-associated protein kinase domain-3 (DAPK-3) is linked with enhanced apoptosis. While coinfection of these two oncogenic pathogens downregulated this, pro-apoptotic which supported the accumulation of AGS gastric epithelial cells over the period. Results showed a significantly lower expression of gankyrin in 12 and 24hrs post-infection in comparison of 48hrs. Furthermore, this finding supported that *H. pylori* and EBV infection mediated higher expression of gankyrin led to aggressive GC through the dysregulation of cell-cycle regulatory genes.

EBV and *H. pylori* can hijack one of the GC markers non-receptor tyrosine kinase (Abl-1) signaling to remodel the host cytoskeleton for multiple purposes such as late phase autophagy, cell motility, and cell adhesion. Trefoil factor-2 (TFF-2) is another gene that is known to be overexpressed in GC. It is a structural component of gastric mucosa and inhibits gut acid secretion and its motility. Inhibition of acid secretion provides the platform for the aggressive growth of *H. pylori* which can further enhance the EBV reactivation and replication (37, 38). Homeobox protein (CDX-2) preferentially binds to methylated DNA and is involved in multiple cellular processes such as transcriptional regulation of genes of epithelial cells. In the present study, profiles of GC marker genes suggested that *H. pylori* changed the status of GC related marker genes for their growth and development of oncogenesis. It was not only involved in the dysregulation of host oncogenic factors but also changed the expression of genes that provided a microenvironment for its growth and replication.

Matrix metalloproteinase-3 and 7 (MMP-3 & MMP-7) is an enzyme that increases the expression of pro-collagenase and degrades the collagen layer, fibronectin, and laminin. The overexpression of this protein in infection models indicates the probable role of EBV and *H. pylori* in gankyrin mediated aggressiveness of GC through the recruitment of such kinds of cell migratory genes.

Downregulated form of the phosphatase and tensin homolog (PTEN) is associated with several types of cancers and functions as a negative regulator of the PI3-kinase/AKT pathway by downregulating PIP3 (phosphatidylinositol 3,4,5-triphosphate) levels. The protein gankyrin downregulated the tumor suppressor protein-16 (p-16), protein-53 (p-53), and protein retinoblastoma (pRB) through the direct interaction with its various domains (17). The loss of activity of these tumor suppressor genes is the key marker for carcinogenesis. This demonstrates that the association of infection of EBV and *H. pylori* in gankyrin upregulation, cellular genome, and dysregulation of cellular signaling was important for oncogenesis.

Apaf-1 and Bax are mitochondrial enzymes that lead to the release of cytochrome c and CASP3 and efficiently initiate the process of apoptosis. Apoptotic protease activating-factor-1 (Apaf-1) and apoptosis regulator (Bax) profiles after coinfection suggested that the coinfection of EBV and *H. pylori* downregulated the proapoptotic genes via upregulating the oncoprotein gankyrin. Furthermore, EBV and *H. pylori* infection mediated upregulated gankyrin downregulates the anti-apoptotic B-cell lymphoma-2 (BCL-2).

The current study demonstrated that EBV and *H. pylori*-associated factors specifically enhanced the expression of gankyrin. Upregulated gankyrin mediated dysregulation of expression of the cell-cycle regulator, GC marker, cell migration, DNA response, and antiapoptotic genes in infected gastric epithelial cells. EBV and *H. pylori* create intracellular and extracellular milieu which potentially favor gastric epithelial cell proliferation and carcinogenesis. It provided the *H. pylori*-EBV tie, which forms a suitable tumor microenvironment for gastric epithelial cell proliferation and carcinogenesis. Further gankyrin might be playing a central role in this transformation process by dysregulation of multiple associated genes **Fig 10**.

**Figure 10:**
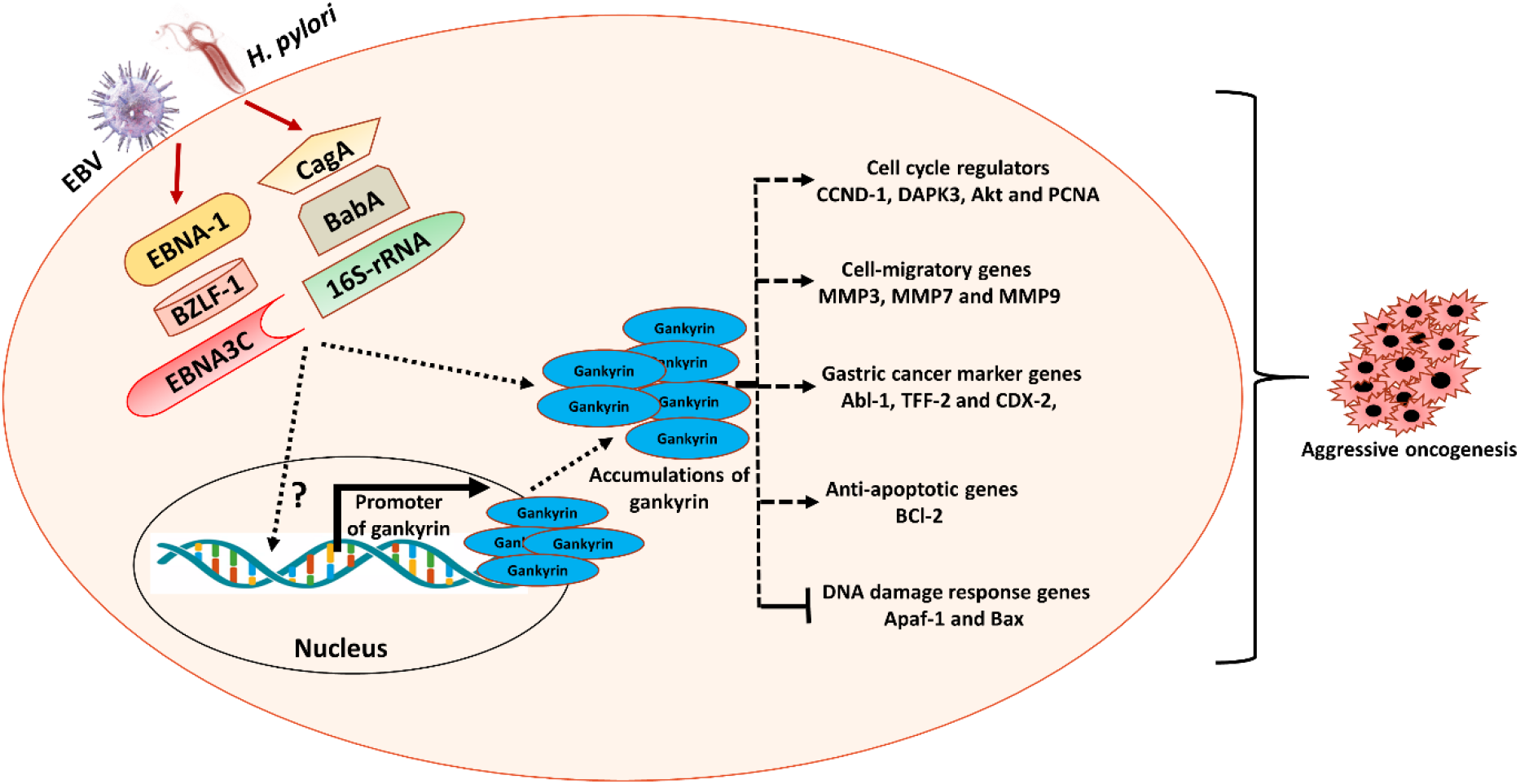
A model illustrating the association of gankyrin in EBV and *H. pylori* mediated gastric cancer. This association drives the progression of EBV and *H. pylori* mediated gastric cell transformation. Oncogenic activity of gankyrin is accelerated in the presence of pathogen which targets the cell cycle regulators, cell migratory genes, gastric cancer marker, anti-apoptotic, DNA damage response and tumor suppressor genes thus caused the aggressive oncogenesis.

## Materials and methods

### Cell culture

The gastric cell lines AGS were obtained from the National Center for Cell Science (NCCS) Pune India. AGS cell line which is an EBV negative cells, cultured in Dulbecco’s modified Eagle’s medium (DMEM; Himedia, Mumbai, India) containing 10% fetal bovine serum (FBS; BIOWEST, South America origin) and 100 U/mL penicillin/streptomycin (Himedia, Mumbai India) under conditions of 5% CO_2_ and humidified air at 37°C (Eppendorf India).

### Virus (EBV) culture and isolation

HEK293T cells containing GFP-EBV (obtained as a gift from the Erle S. Robertson’s laboratory, University of Pennsylvania, USA) were cultured and used for EBV amplification. Cultured the virus and purified as described previously (2, 39, 25)

### Bacterial (*H. pylori*) culture

*H. pylori* strain (I10) was a kind gift by Dr. Asish Kumar Mukhopadhyay (National Institute of Cholera and Enteric Diseases, ICMR, Kolkata, India) *H. pylori* was grown according to the conditions described earlier (25).

### Coinfection of EBV and *H. pylori* with AGS cells

For infection, we took 0.25 million AGS cells and cultured in a 6-well plate. Observed the cells up to the 40-50% confluencies and then incubated the MOI-100 of *H. pylori* and 25μl of EBV. We optimized the EBV dose by incubating the increasing dose of EBV such 25, 50, 100, 150, and 200 μl with AGS cells and observed 25 μl of EBV culture was sufficient for a successful infection. For this study, we have developed five independent approaches for the construction of coinfection models by using *H. pylori* and EBV. In the first approach, uninfected AGS cells were taken as the negative control (**Fig.1A** **control)**. In our second and third approaches, we have incubated the *H. pylori* and EBV with AGS cells (**Fig.1B** **infection-I and C Infection-II**) respectively. Moreover, in the fourth approach, we have incubated the *H. pylori* with AGS cells for six hours thereafter given the infection of EBV and incubated for 12, 24, and 48 hours (**Fig.1D** **infection-III**). In the fifth approach, the infection of EBV is given first for six hours followed by the infection of *H. pylori* (**Fig.1E** **Infection-IV**).

### RNA isolation and qRT-PCR

Fixed (MOI-100) of *H. pylori* (I10) and EBV (25μl) were incubated with AGS cells under specific conditions (5% CO_2_, 37 °C) for 12, 24, and 48 hrs. RNA isolated and qRT-PCR was executed as described earlier (25). The pathogens and host genes were analyzed.

### Western Blot

Western blot experiments were carried out as described earlier (40). The lysates were analyzed by Western blots using an anti-gankyrin polyclonal primary antibody (cell signaling) and HRP-tagged secondary antibody. Blots were observed under the gel doc system (ChemiDoc™ XRS+ System with Image Lab™ Software #1708265)

### Immunofluorescence Assay

An immunofluorescence assay was performed as described earlier (40) by using an anti-gankyrin primary antibody and specific signals were detected with secondary antibodies conjugated with Alexa Fluor 488 (cell signaling). The cell’s nucleus was counterstained with 4′, 6′-diamidino-2-phenylindole (DAPI). The results were analyzed with confocal microscopy (Olympus IX83) at 60X and zoom two times in triplicates.

### Cell proliferation assay

An assay of cell proliferation was accomplished through trypan blue exclusion and crystal violet staining methods as described earlier (2, 40).

### Colony formation assay

The AGS cells were infected with EBV and *H. pylori* and EBV positive cells selected with puromycin (2ug/ml). We selected the ectopically expressed gankyrin positive AGS cells through geneticin (G148). The image was taken by using ChemiDoc™ XRS+ System with Image Lab™ Software #1708265 (40).

### Statistical Analysis

Data were statistically analyzed using a two-tailed Student’s *t*-test. All the results were derived from triplicate experiments P-values by using the Graph Pad Prism version 6 of <0.05, <0.01, and <0.001 were considered statistically significant and represented with *, ** and ***, denoting downregulation/upregulation respectively.

## Acknowledgements

This project was supported by the Council of Scientific and Industrial Research grant number 37(1693)/17/EMR-II and Department of Science and Technology as Ramanujan fellowship grant no. SB/S2/RJN-132/20/5. We are thankful to CSIR, UGC and DST-Inspire for fellowship to Dharmendra Kashyap, Budhadev Baral, Nidhi Varshney respectively in the form of research stipend. We appreciate Dr. Asish Kumar Mukhopadhyay (National Institute of Cholera and Enteric Diseases, Kolkata) for providing the *Helicobacter pylori* strain I10. We are thankful to Dr. Erle S Robertson (University of Pennsylvania, USA) for providing us with HEK293T EBV BAC cell, which consistently expressed Epstein Barr Virus (EBV) genome. We also appreciate the Sophisticated Instrumentation Centre, IIT Indore for confocal microscopy facility. We gratefully acknowledge the Indian Institute of Technology Indore for providing facilities and support. We appreciate Ms. Shweta Jakhmola and our other lab colleagues for insightful discussions and advice. We are thankful to Mr. M Israel and Mr. Prajwal M for their voluntary help in some experiments.

## Abbreviation

*H. pylori*: *Helicobacter pylori*
EBV: Epstein Barr Virus
GC: Gastric cancer
EBNA1: Epstein Barr Nuclear antigen-1
EBNA3C: Epstein Barr Nuclear antigen-3C
EBNA2: Epstein Barr Nuclear antigen-2
BZLF-1: BamHI Z fragment leftward open reading frame 1
16s rRNA: 16s Ribosomal RNA
CagA: cytotoxin-associated gene A
BabA: Blood group antigen binding adhesin A
VacA: Vacuolating toxin A
pRb: protein retinoblastoma
P53: Tumor suppressor protein 53
AGS: Adenogastric carcinoma cell
CCND-1: Cyclin D1
DAPK3: Death associated protein kinase domain-3
PCNA: Proliferating cell nuclear antigen
Akt: Protein kinase B
PTEN: Phosphatase and tensin homolog
APC: Adenomatous polyposis coli
Abl-1: Tyrosine-protein kinase ABL1
TFF2: Trefoil factor-2
CDX-2: Homeobox protein CDX-2
MMP-3: Matrix metalloproteinase-3
MMP-7: Matrix metalloproteinase-7
Apaf-1: Apoptotic protease activating-factor-1
Bax: Apoptosis regulator BAX
Bcl-2: B-cell lymphoma-2
CCL8: Chemokine ligand 8
CagA-PAI: CagA-pathogenicity island
PIP3: phosphatidylinositol 3,4,5-triphosphate

## Supplementary Figure and figure legend

**Supplementary Fig.1.**
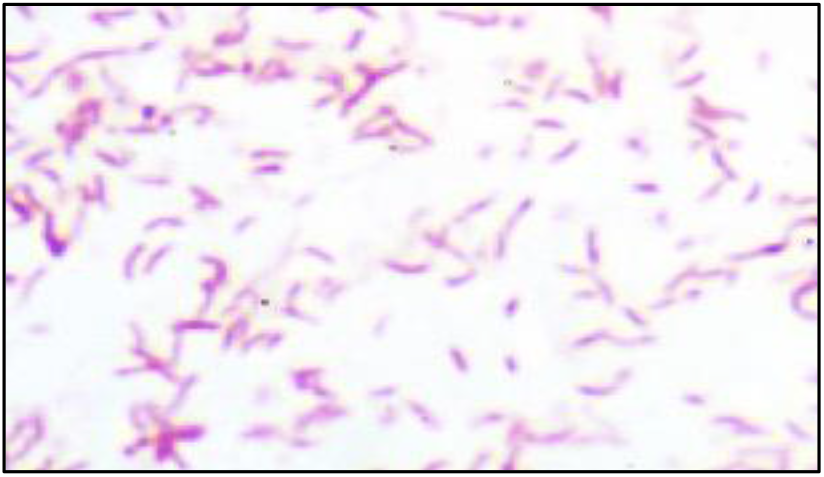
Gram staining of *H. pylori* (I10) showing the regular spiral shape of bacteria

